# SpatialCells: Automated Profiling of Tumor Microenvironments with Spatially Resolved Multiplexed Single-Cell Data

**DOI:** 10.1101/2023.11.10.566378

**Authors:** Guihong Wan, Zoltan Maliga, Boshen Yan, Tuulia Vallius, Yingxiao Shi, Sara Khattab, Crystal Chang, Ajit J. Nirmal, Kun-Hsing Yu, David Liu, Christine G. Lian, Mia S. DeSimone, Peter K. Sorger, Yevgeniy R. Semenov

## Abstract

**Background:** Cancer is a complex cellular ecosystem where malignant cells coexist and interact with immune, stromal, and other cells within the tumor microenvironment. Recent technological advancements in spatially resolved multiplexed imaging at single-cell resolution have led to the generation of large-scale and high-dimensional datasets from biological specimens. This underscores the necessity for automated methodologies that can effectively characterize the molecular, cellular, and spatial properties of tumor microenvironments for various malignancies.

**Results:** This study introduces SpatialCells, an open-source software package designed for region-based exploratory analysis and comprehensive characterization of tumor microenvironments using multiplexed single-cell data.

**Conclusions:** SpatialCells efficiently streamlines the automated extraction of features from multiplexed single-cell data and can process samples containing millions of cells. Thus, SpatialCells facilitates subsequent association analyses and machine learning predictions, making it an essential tool in advancing our understanding of tumor growth, invasion, and metastasis.

**Availability of code and materials:** https://github.com/SemenovLab/SpatialCells.

## Introduction

Cancer presents an intricate cellular ecosystem, which plays a critical role in tumor development, progression, and therapeutic outcomes. The spatial organization of cells and their interactions within the tumor microenvironment (TME) contain essential insights into the course of tumor growth and progression. Characterizing molecular, cellular, and spatial properties of TMEs across diverse malignancies has gained substantial attention^1,2^.

Recent technological breakthroughs in spatially resolved multiplexed imaging at single-cell resolution, such as CODEX^3^ and CyCIF^4^, are highly effective in studying TMEs and intratumoral heterogeneity within solid tumors^5,6^. However, the lack of systematic computational methodologies to leverage the large volume of data generated by these technologies poses a significant challenge for their scalable deployment in clinical settings.

Currently, the available tools for handling multiplexed imaging data are designed to address specific aspects of multiplexed imaging data analysis. Some tools focus on converting multi-channel whole-slide images into single-cell data (e.g., MCMICRO^7^), while others prioritize preprocessing, cell type phenotyping, and visualizing the obtained single-cell data (e.g., SCIMAP^8^). Spatial analysis of single-cell data represents only a small portion of the functions in existing packages, such as HALO^9^, and often demands manual annotation. Consequently, the capability of these tools to effectively conduct comprehensive spatial analysis remains limited.

Furthermore, the considerable progress in leveraging machine learning for predictive tasks presents a critical consideration for clinical outcomes. However, these algorithms often require substantial sample sizes to achieve robust performance. Overall, the high-dimensional, heterogeneous, and complicated dependency structures inherent in multiplexed single-cell data, coupled with the need for large sample sizes in machine learning algorithms, present challenges to conventional manual annotation and statistical techniques. Thus, there is an urgent need to develop computational methodologies to analyze multiplexed single-cell data in a scalable manner and enable the development of forecasting models that would inform clinical decision-making and enhance our understanding of disease progression.

In this study, we introduce SpatialCells, an open-source software package designed to perform region-based exploratory analysis and characterization of TMEs using multiplexed single-cell data. This tool is equipped to efficiently analyze tissue samples containing millions of cells and automatically extract quantitative features, enabling subsequent association analyses and machine learning predictions at scale.

## Implementation

### Overview

This study is grounded in existing literature on tumor and immune parameters associated with cancer progression and patient survival. Our primary goal is to develop automated methodologies for quantifying these parameters with spatially resolved single-cell data. By integrating clinicopathologic features extracted from electronic medical records, we will be equipped to perform a comprehensive association analysis or make predictions about patient outcomes, as illustrated in Figure 1.A and Figure 1.B. We have provided detailed tutorials on data exploration, tumor cell-centric analysis, and immune cell-oriented analysis at https://github.com/SemenovLab/SpatialCells.

**Figure 1.**
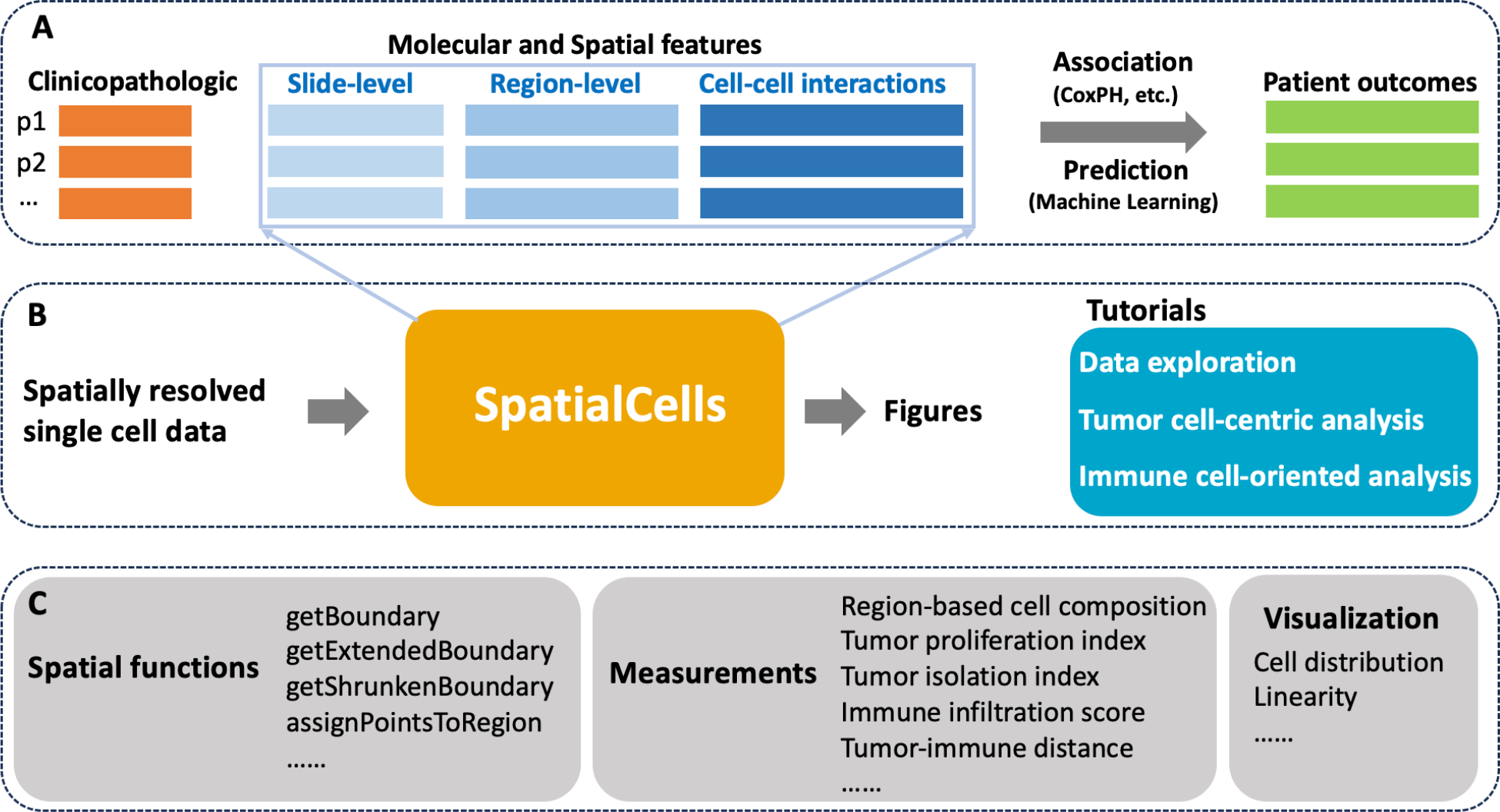
Overview of the framework. **A**. The role of SpatialCells within downstream analyses. In this context, “p1, p2, …” represent patient IDs, and CoxPH (Cox Proportional-Hazards Modeling) is one of the association analysis methods. **B**. The input and output of SpatialCells: It takes the spatially resolved multiplexed single cell data as input and provides quantified features along with the corresponding visualizations as output. Tutorials for data exploration, tumor cell-centric analysis, and immune cell-oriented analysis have been provided. **C**. Key modules in SpatialCells: SpatialCells incorporates several main modules to facilitate its functionality, such as tumor proliferation index, immune infiltration score, and tumor-immune distance.

SpatialCells is featured by its capability to define regions of interest based on any group of cells and subsequently conduct region-based analyses. Figure 1.C presents the main modules incorporated into SpatialCells. Our workflow starts with developing a *Spatial* module, including functions to establish regional boundaries and check the region in which a cell is located. The *Measurements* module contains functions to extract the properties of tumor cells and tumor-immune cell interactions, including tumor proliferation index, immune infiltration score, and tumor-immune cell distances. These properties can be assessed for the whole tissue or local regions. Importantly, our methods can efficiently process datasets containing millions of cells.

### Data exploration

In the data exploration, we have incorporated measurements and visualizations that can provide initial insights into the samples. These include:

- **Whole-slide level cell composition**. This involves assessing the percentages of different cell types, such as tumor cells and immune cells, present in the samples.
- **Region-of-interest level cell composition**. Whole-slide tissues often have a significant amount of blank background. SpatialCells computationally defines regions of interest (ROIs) and removes background in a standardized manner across all tissues in the study cohort. This is accomplished by initially calculating the tumor boundary and then extending this boundary by a specified distance (a user-defined parameter).
- **Region-based cell composition**. SpatialCells provides the capability for geometric partition of tissues in various ways. For instance, it can divide the tumor region into subregions based on distance from the centroid or angle from the zero degree. Following the partitioning, SpatialCells can enumerate cell types within each tissue subregion for detailed compositional analysis.
- **Region-based clustering**. Similarly, after partitioning, SpatialCells provides the ability to cluster cells within a specific region.

### Tumor cell-centric analysis

Tumor cell-centric analysis focuses on the characterization and study of tumor cells to gain insights into their biology, heterogeneity, and behavior within the TME.

Specifically, the American Joint Committee on Cancer/Union for International Cancer Control (AJCC/UICC) staging system provides guidelines for classifying the extent of cancer spread, considering factors such as tumor size and depth of invasion^10–12^. Additionally, the mitotic rate of various tumors is highly relevant in predicting patient outcomes^13–15^. In this analysis, we have emphasized the following tumor characteristics that may be linked to patient prognosis:

- **Tumor area and tumor cell density**. SpatialCells offers functions for defining regions based on user-specified markers and calculating the area of a region and the density of any cell type within the region.
- **Tumor multivariate proliferation index**. Gaglia et al. demonstrated the effectiveness of a multivariate proliferation index (MPI) to differentiate proliferating from non-proliferating tumor cells^16^. Building on this work, SpatialCells provides a function for calculating a multivariate proliferation index of a given cell type, including tumor cells. The input of this function includes two marker lists of interest: 1) mitosis/proliferation markers (e.g., phosphohistone H3, Ki67, PCNA, and MCM2); and 2) cell cycle arrest markers (e.g., p21 and p27). Adapted from the definition in the work by Gaglia et al^16^, the MPI is defined as follows:

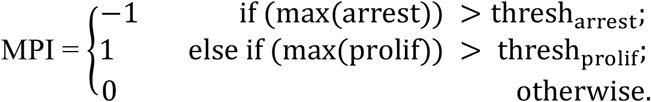

The threshold values for proliferation and arrest are data dependent. For normalized markers of expression levels from 0 to 1, the values are set to 0.5 by default. The thresh_arrest_ can be tuned on the basis of a commonly used proliferation marker, such as Ki67. Instead of computing a single MPI for the whole tissue, SpatialCells enables calculating MPI for subregions of interest within the tissue sample. We demonstrate this method by applying a sliding window of a user-defined size over the tissue and computing the MPI for each subregion.

- **Tumor isolation index**. Categorizing tumors into immune hot, immune suppressed, and immune cold/isolated groups has considerable prognostic value in various malignancies^17,18^. Here we focus on defining a tumor isolation index, which is the fraction of tumor cells in the region without immune cells over all tumor cells. This index is accomplished by dividing the overall region of interest (whole-slide tissue with background removed) into two subregions: immune-rich region and region with almost no tumor-infiltrating immune cells.

### Immune cell-oriented analysis

The TME is a spatially organized landscape, characterized by the presence of lymphocytes, macrophages, and other cell types located both within the central region and at the invasive margin of the tumor. Understanding the TME composition is essential for developing effective cancer therapies and predicting patient prognosis. Here, we have conducted macro-region and micro-region analysis, along with cell-cell interactions:

- **Macro-region analysis**. The overall region of interest, which encompasses the whole-slide tissue with the background removed, is divided into four subregions: tumor, tumor border, stroma border, and stroma, as illustrated in Figure 2.A. We subsequently evaluate the cell compositions within each region, with a particular focus on immune cells.

**Figure 2.**
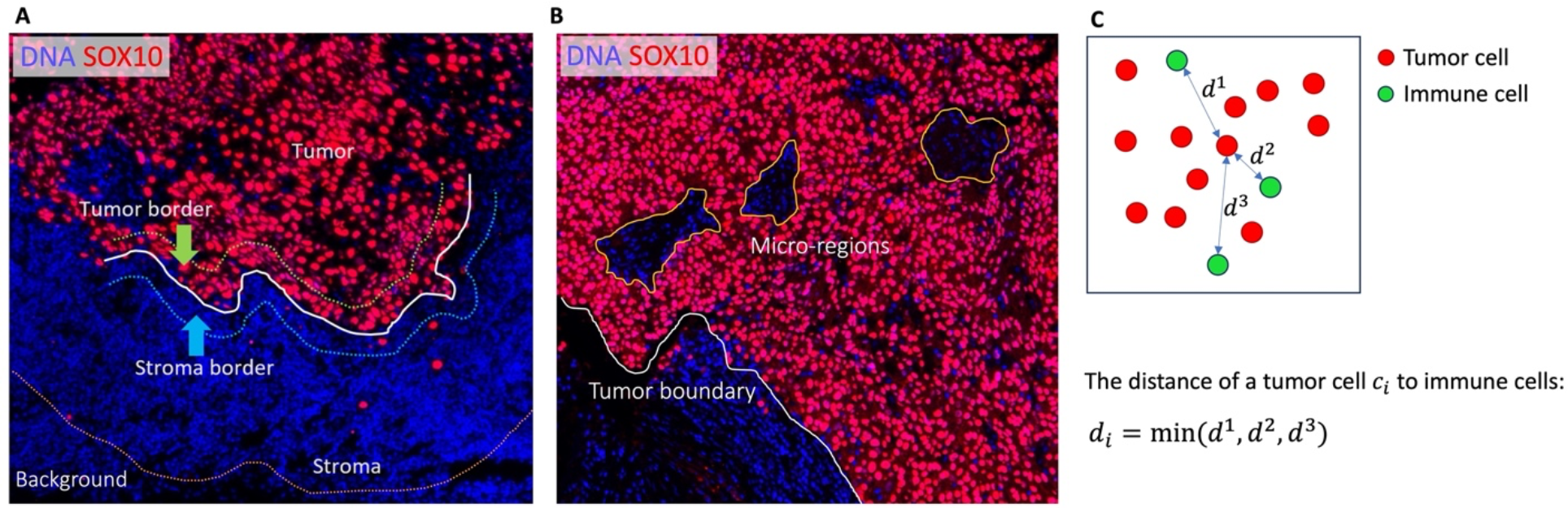
Macro-regions, micro-regions, and cell-to-cell distance. **A**. The background defined based on a user-specified distance from the tumor boundary is removed from the whole-slide image. The remaining region of interest is partitioned into tumor region, tumor border, stroma border, and stroma region. **B**. Within the tumor region, there are small regions of interest, defined as micro-regions, which allow fine-grained analyses. **C**. The distance of a cell (e.g., a tumor cell) to another cell type (e.g., immune cells) is defined as the distance between the cell and its nearest neighbor of another cell type.
- **Micro-region analysis**. This analysis focuses on small regions, defined as micro-regions, within the tumor area, as illustrated in Figure 2.B. By identifying these micro-regions, we can perform detailed characterizations of cells within them, enabling a finer-grained understanding of the cellular landscape.
- **Cell-cell interactions**. An important aspect of our analysis involves investigating the interactions between various cell types within the TME. For instance, SpatialCells can be applied to quantify the interactions between tumor cells and immune cells. As shown in Figure 2.C., we assess the degree of tumor immune infiltration by measuring the distance between a tumor cell and its nearest immune cell. This information provides valuable insights into the interplay between different cell populations within the TME.

## Results

In this section, we demonstrate the functionality of SpatialCells by presenting results in data exploration, tumor cell-centric analysis, and immune cell-oriented analysis. Additional analyses can be found in tutorials, providing researchers with a comprehensive toolkit for in-depth investigations.

Our analyses involve a publicly available multiplexed imaging data of a cutaneous melanoma sample (MEL1) consisting of 1,110,585 cells^19^. MEL1 was imaged using CyCIF^4^ with 30 antibody markers (e.g., SOX10, Keratin, CD31, CD3D, CD8A, CD11C, MITF, and Ki67) and preprocessed using MCMICRO^7^, which transforms multi-channel whole-slide images into single-cell data. The output of MCMICRO is the input of SpatialCells. Additional details of this sample can be found in the provided reference^19^.

### Data exploration

SpatialCells is a versatile tool for region-based data exploration, particularly for assessing cell composition or clustering within whole-slide images or specific regions of interest. In Figure 3, we illustrate how SpatialCells can be applied to compute cell compositions for three different types of regions within the MEL1 sample. We focus on the main tumor area for clarity.

**Figure 3.**
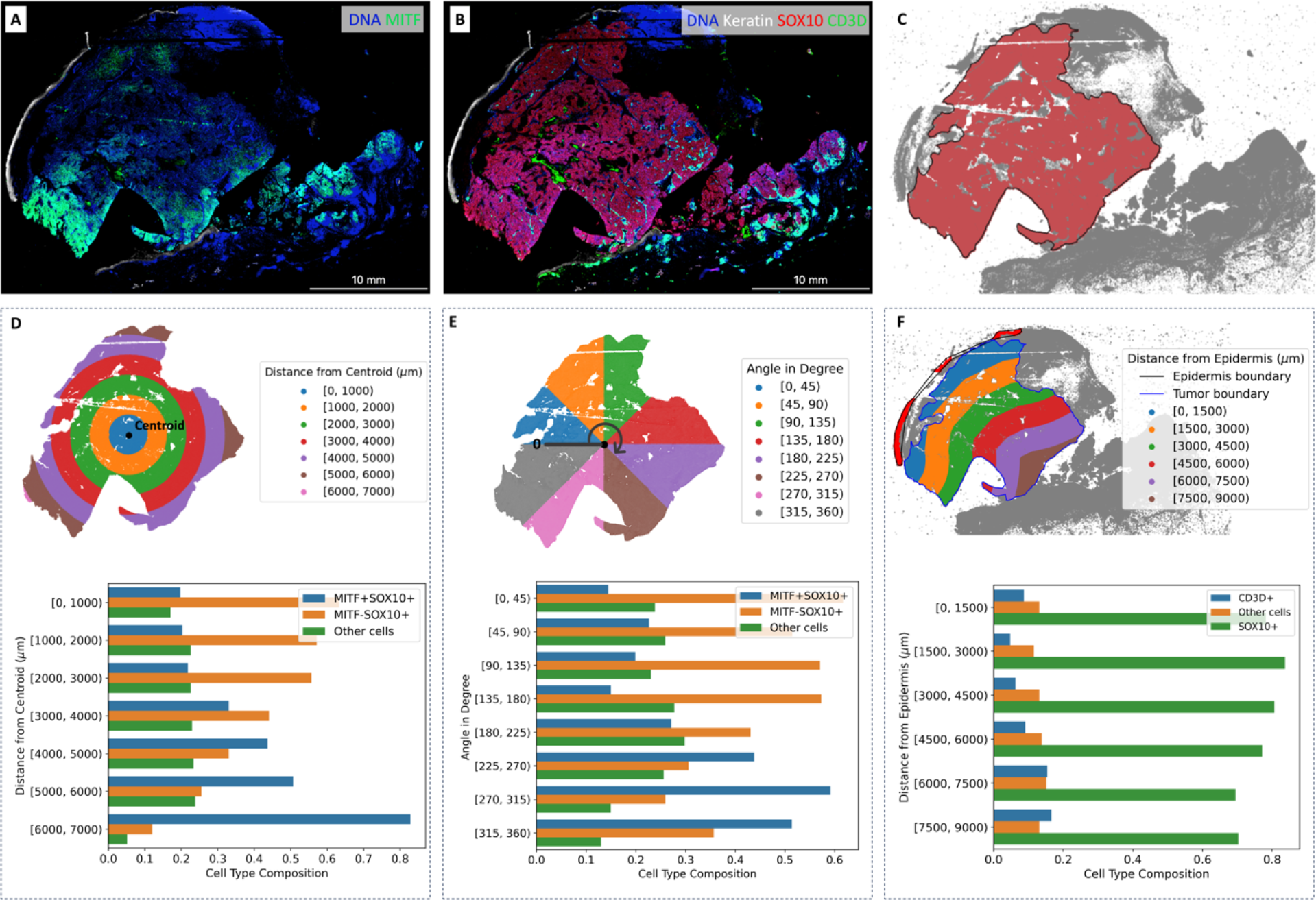
Cell composition in regions of interest. **A**. Gradient of MITF on the CyCIF imaging data. **B**. Keratin, SOX10, and CD3D expressions on the CyCIF imaging data. **C**. Boundary of the main tumor area. **D**. Cell type composition in subregions based on distance from the centroid. **E**. Cell type composition in subregions based on the angle from the zero-degree reference. **F**. Cell type composition in subregions based on the distance from the epidermis.

Figure 3.A and Figure 3.B present the expression levels of related markers, including Keratin, SOX10, MITF, and CD3D, within the imaging data. SOX10 expression is a sensitive and specific marker for melanoma tumor cells. MITF is an important melanoma oncogene and plays a role in determining therapeutic resistance^20,21^. CD3D is a marker commonly associated with immune cells, particularly T lymphocytes (or T cells). Figure 3.C shows the boundary of the main tumor area as identified by SpatialCells. In Figure 3.D, the subregions are defined based on the distance from the centroid of the main tumor area. In Figure 3.E, the subregions are based on their angles from the zero-degree reference. The percentage of MITF+SOX10+ cells within each subregion is computed. The two barplots in Figure 3.D and Figure 3.E demonstrate the gradient of MITF+ tumor cells. Figure 3.F provides cell composition within subregions, which are defined based on their distance from the epidermis. Both the centroid (start-point) and the epidermis (start-line) are computationally determined. It is worth noting that SpatialCells offers support for user-defined start-points and start-lines, allowing users to tailor analyses that align with specific requirements.

### Tumor isolation index and multivariate proliferation index

SpatialCells also facilitates tumor-cell-centric analyses. Figure 4 showcases the tumor isolation index and the MPI. These analyses are part of our dedicated tumor-cell-centric tutorial.

**Figure 4.**
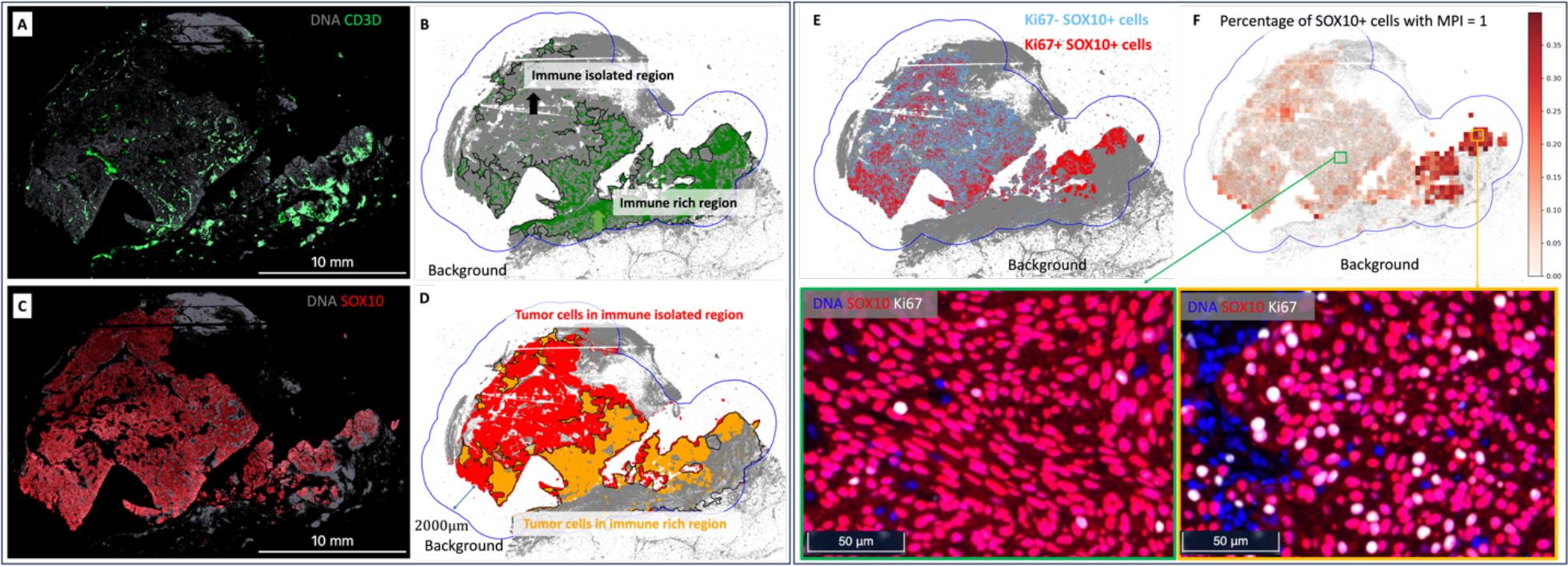
Tumor isolation index (**A, B, C**, and **D**) and tumor multivariate proliferation index (**E** and **F**). **A**. Expression level of CD3D+ cells in the MEL1 sample. **B**. Division of the overall ROI into two subregions based on CD3D+ cells: 1) immune-isolated region, which has almost zero immune cells; 2) immune-rich region. **C**. Expression level of tumor (SOX10+) cells in the MEL1 sample. **D**. Visualization of tumor cells in the immune-isolated region and immune-rich region, respectively. The tumor isolation index for this sample is 46.9%, which quantifies the percentage of tumor cells in the immune-isolated region. **E**. Display of Ki67+ and Ki67-tumor cells. Ki67 is a proliferative marker. The overall percentage of proliferative tumor cells (MPI = 1) in the MEL1 sample is 7.4%. **F**. Percentage of tumor cells with MPI = 1 in subregions and the zoomed-in CyCIF images. A sliding window of size 300 × 300 microns is applied over the ROI to compute the percentage of tumor cells with MPI = 1. In this example, the MPI is computed using Ki67 due to the availability of markers. Additionally, zoomed-in CyCIF images are provided to visualize areas with varying levels of Ki67+ tumor cells.

Figure 4.A shows the expression of CD3D+ cells in the MEL1 sample. In Figure 4.B, the overall ROI is divided into two distinct subregions based on the presence of CD3D+ cells: 1) immune-isolated region, characterized by minimal immune cell presence; 2) immune-rich region, characterized by a higher presence of immune cells. Figure 4.C shows the expression of SOX10+ cells in the MEL1 sample. Figure 4.D visually contrasts tumor cells in the immune-isolated region and the immune-rich region. The tumor isolation index for this sample is quantified as 46.9%, which is the percentage of tumor cells in the immune-isolated region.

Figure 4.E. displays Ki67+ and Ki67-tumor cells. Ki67 is a marker of proliferation. The overall percentage of proliferating tumor cells (MPI = 1) in the MEL1 sample is 7.4%. Figure 4.F shows the percentage of tumor cells with MPI = 1 in a sliding window of 300 × 300 microns. In this example, the MPI is computed using Ki67 due to the availability of markers. Additionally, zoomed-in CyCIF images are provided to visualize areas with varying levels of Ki67+ tumor cells.

### Macro-region and micro-region analyses

The immune context within the TME plays an important role in cancer prognosis and therapeutic efficacy^18^. Figure 5 provides a comprehensive overview of both macro-region (A, B, and C) and micro-region (D, E, F, G, and H) analyses, empowering users to comprehensively characterize the immune and cellular landscape within the TME.

**Figure 5.**
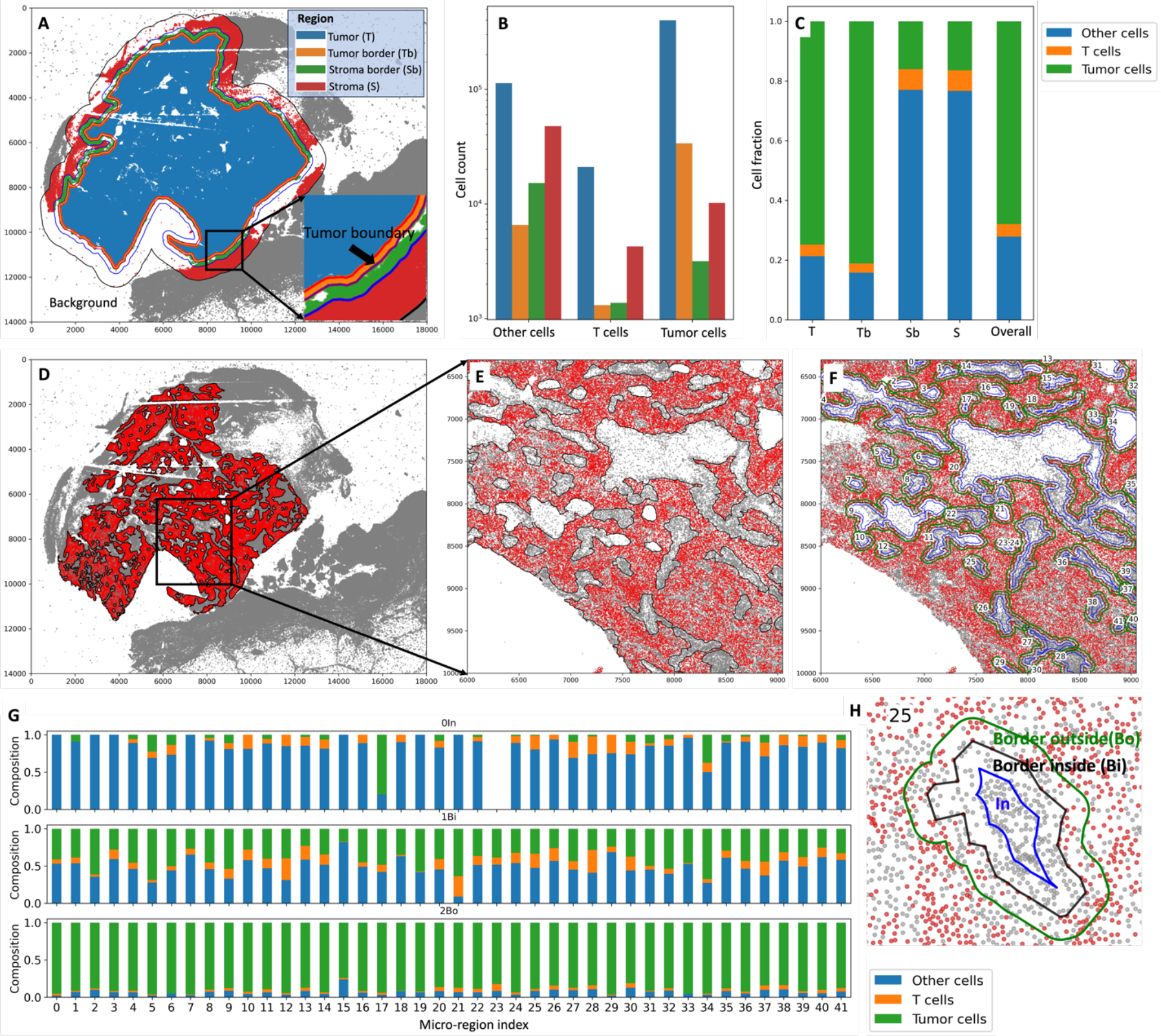
Macro-region analysis (**A, B**, and **C**) and micro-region analysis (**D, E, F, G**, and **H**). **A**. The overall region of interest is divided into four subregions: Tumor (T), Tumor border (Tb), Stroma border (Sb), and Stroma (S). The tumor region boundary is 100 microns away from the tumor boundary; the stroma border region boundary is 200 microns away from the tumor boundary; the boundary for the overall region of interest is 800 microns away from the tumor boundary. Users can specify these distances based on their needs. **B**. The corresponding counts of tumor cells, T cells, and other cells in the four regions. **C**. The cell fraction among all cells within each region. **D**. The micro regions within the tumor area. **E**. The zoomed-in micro regions. Each micro-region is indexed, allowing users to pick the ones of interest. **F**. Each micro-region boundary is extended and shrank (offset = 30 microns) to get the “In”, “Border inside”, and “Border outside” subregions, as shown in **H. G**. Cell compositions within the three subregions for each micro-region.

In Figure 5.A, the overall region of interest is divided into four distinct subregions: Tumor (T), which represents the core tumor region; Tumor border (Tb), which is the transitional area immediately adjacent to the tumor; Stroma border (Sb), marking the border between tumor and stroma; and Stroma (S), situated farther from the tumor. These subregions are defined based on specific distances from the tumor boundary, with the tumor region boundary set at 100 microns, the stroma border region boundary at 200 microns, and the boundary for the overall region of interest at 800 microns away from the tumor boundary. Importantly, users have the flexibility to customize these distances. In Figure 5.B, the corresponding counts of tumor cells, T cells, and other cells are presented for each of the four defined regions. Figure 5.C further provides insights by illustrating the cell fraction among all cells within each of the defined regions. Figure 5.D displays micro-regions within the tumor area. Each micro-region is indexed, enabling users to pinpoint specific micro-regions of interest. Figure 5.E offers a closer look at these micro-regions. In Figure 5.F, every micro-region boundary is extended and contracted (offset = 30 microns) to define “In”, “Border inside”, and “Border outside” subregions, as visually demonstrated in Figure 5.H. The cell compositions within these three subregions for each micro-region are presented in Figure 5.G.

## Discussion and future directions

SpatialCells represents a significant advance in the field of spatial analysis for multiplexed single-cell image data, enabling the computational quantification and standardized analyses of critical features within the TME. Its functionalities encompass various aspects of TME analysis, such as region-based cell composition, tumor proliferation index, tumor isolation index, immune cell infiltration, and tumor-immune distance. By providing these analytical tools, SpatialCells empowers researchers to gain deeper insights into the intricate spatial relationships and characteristics of cells within the TME.

One of the limitations of the SpatialCells software is that the cell segmentation provided by upstream imaging data preprocessing tools, such as MCMICRO^7^, may introduce errors in the cell counts. However, SpatialCells is a customizable software that can analyze multiple samples within a study in a consistent and standardized manner. This capability involves applying similar settings and parameters across all samples, thus mitigating some of the variability introduced by cell segmentation. Another limitation of the SpatialCells software is the reliance on the quality of gating and calling cell types, which involve converting continuous marker expression levels into binary variables. This binary conversion introduces subjectivity, leading to variability in cell count. Finally, the adaptability of SpatialCells, with its support for user-defined parameters, affords users the freedom to configure settings based on their specific needs. However, this flexibility can also be a source of errors, as suboptimal parameter choices may impact the accuracy of the analysis.

In the future, we will continue to improve this software and provide clear guidelines to users on best practices for parameter selection. We will include more functions as well as tutorials, particularly for multimodal analyses within the field of spatial biology. For example, we will provide functions for migrating ROIs from other modalities, such as Hematoxylin and Eosin (H&E) stain imaging and spatial transcriptomics, to multiplexed imaging, which will allow integrative analyses across different modality data.

## Conclusions

In summary, SpatialCells is a novel software solution for spatially analyzing the TME using multiplexed imaging data in a streamlined fashion with the capacity to process samples containing millions of cells. This software is of critical importance in the analysis of the TME and in furthering our understanding of the factors leading to tumor progression as it facilitates subsequent association analyses and machine learning predictions.

## List of abbreviations

TME: Tumor Microenvironment
CODEX: Co-detection by indexing: a method for multiplexed tissue imaging
CyCIF: Cyclic immunofluorescence: a method for highly multiplexed tissue imaging
SCIMAP: A toolkit for analyzing spatial molecular data.
HALO: A toolkit for quantitative image analysis
ROI: Region of interest
AJCC: American Joint Committee on Cancer
UICC: Union for International Cancer Control
MPI: Multivariate proliferation index

## Declarations

### Authors’ contributions

All co‐authors provided an intellectual contribution to this study. GW, ZM, BY, TV, YS, PKS and YRS conceived and designed the study. GW and BY wrote the scripts, implemented the SpatialCells package, and wrote the tutorials. GW, BY, SK, and ZM generated the figures and wrote the manuscript. The study was conducted under the supervision of PKS and YRS. All authors reviewed and approved the manuscript.

